# The CD56bright CD62L+ NKG2A+ immature cell subset is dominantly expanded in human cytokine-induced memory-like NK cells

**DOI:** 10.1101/405134

**Authors:** Juhyun Jin, Yong-Oon Ahn, Tae Min Kim, Bhumsuk Keam, Dong-Wan Kim, Dae Seog Heo

## Abstract

Recent studies have revealed immunological memory of NK cells. Short-term *in vitro* cytokine stimulation also induces NK cell memory, but heterogeneous cell subsets within the cytokine-induced memory-like (CIML) NK cells has not been elucidated. Here we found that the dominant cell subset in human CIML NK cells are immature CD56^*bright*^ CD62L^+^cells, and they were selectively expanded CD56^*bright*^ CD16^*-*^ CD62L^+^ NK cells. Although these cells acquired KIR expression after the cytokine stimulation, sustained NKG2A expression inhibits cytotoxicity against HLA-E^+^ target cells. In contrast, another checkpoint molecule LAG-3 is induced mainly on KIR^+^ NKG2C^+^ minor CIML NK cells. Our findings imply targeting NKG2A and LAG-3 should be considered for CIML NK cell-based immunotherapy.

## Introduction

Natural killer (NK) cells are large granular lymphocytes of the innate immune system that do not have antigen specificity unlike T or B lymphocytes (Vivier, Tomasello, Baratin, Walzer, & Ugolini, 2008). Instead, NK cells recognize and eradicate transformed or viral infected cells by detecting stress-induced ligands (Caligiuri, 2008; Ljunggren & Karre, 1990). Although NK cells do not have antigen specificity, recent studies have shown that NK cells also have immunological memory. Murine and human cytomegalovirus (CMV) infection induces immunological memory in a subset of NK cells *in vivo* (Cerwenka & Lanier, 2016; Lee et al., 2015; O’Sullivan, Sun, & Lanier, 2015; Schlums et al., 2015). As these memory NK cells show a unique gene expression profile, these memory NK cells are also called ‘adaptive’ NK cells.

*In vitro* cytokine stimulation also let NK cells have a memory-like function, and these cells are called ‘cytokine-induced memory-like (CIML)’ NK cells. Once *ex vivo* activated murine NK cells by exogenous IL-12 and IL-18 in the presence of low dose IL-15 as a survival factor, had long *in vivo* lifespan when injected into mice, and express abundant IFN-*γ* by cytokine re-stimulation (Cooper et al., 2009). Similar findings were also shown in human NK cells (Leong et al., 2014; Romee et al., 2012), use of adaptive transfer of CIML NK cells are now under clinical trials (Romee et al., 2016). However, CIML NK cells much differ from CMV infection-induced adoptive NK cells in terms of phenotype and function. Adaptive NK cells are terminally maturated NKG2C^+^ CD57^+^ cells with down-regulation of the immunoreceptor tyrosine-based activation motif (ITAM)-bearing adaptor proteins like Fc*ε*RI*γ*, SYK, EAT2, and transcription factor pro-myelocytic leukemia zinc finger (PLZF; also known as ZBTB16) (Lee et al., 2015; Schlums et al., 2015). However, these molecules are not down-regulated in the CIML NK cells. In addition, cytokine responsibility is higher in CIML NK cells while reduced cytokine receptor expression is shown in adaptive NK cells (Leong et al., 2014; Romee et al., 2012).

In human peripheral blood, there are two NK cell subsets which are CD56^*dim*^ CD16^+^ major (about 90%) and CD56^*bright*^ CD16^*-*^ minor cell subsets (Caligiuri, 2008). In contrast to mature, cytotoxic CD56^*dim*^ NK cells, immature CD56^*bright*^ NK cells have lower cytotoxicity. Instead, CD56^*bright*^ NK cells express abundant IFN-*γ* in response to cytokine stimulation. Interestingly, the CIML and CD56^*bright*^ NK cells share several surface markers, such as the presence of NKG2A and the absence or low frequencies of NKG2C, CD57, and killer cell immunoglobulin-like receptors (KIRs) (Romee et al., 2016). CD62L (L-selectin), a member of selectin family, is a cell surface adhesion molecule and a homing receptor that makes lymphocytes enter to secondary lymphoid tissues (Chen, Engel, & Tedder, 1995). CD62L and CD57 are reversely and mutually exclusively expressed in NK cells during their maturation (Bjorkstrom et al., 2010; Lopez-Verges et al., 2010). While CD56^*bright*^ NK cells are CD62L^+^ CD57^*-*^, when the cells are culture with cytokine such as IL-2 or IL-15 *in vitro*, these cells loss CD62L and acquire CD57 expression. Thus, NK cells loss CD62L and acquire CD57 during their differentiation, education, and maturation *in vivo* and *in vitro* (Moretta, 2010). Although less cytotoxic CD56^*bright*^ NK cells also gain cytotoxicity after cytokine activation (Wagner et al., 2017), then differentiate into CD56^*dim*^ NK cells (Romagnani et al., 2007), the relationships of cytokine-activated CD56^*bright*^ and CIML NK cells have not determined yet. In this study, we hypothesized that CD56^*bright*^ NK cells are selectively expanded by cytokine stimulation, and becomes dominant population in CIML NK cells, as these two subsets share same surface markers and functions (enhanced IFN-*γ* secretion by cytokine stimulation).

## Results

### Cytokine-induced memory-like NK cells have immature phenotype

CMV infection-induced adaptive NK cells can be distinguished from the other NK cells by unique surface markers (NKG2C/CD94^+^, CD57^+^, and self KIR^+^), expression of a transcription factor PLZF, and down-regulation of signal transduction molecules (SYK, EAT-2, and/or Fc*ε*RI*γ*) (Lee et al., 2015; Schlums et al., 2015). Although there are several heterogeneous subsets in NK cells, but no known CIML NK cell-specific markers for the dominant subsets. Thus, we first investigated to find the subsets within CIML NK cells. CIML NK cells were generated from peripheral blood lymphocytes from healthy donors according to the standard protocol (supplementary figure Fig. S1) (Romee et al., 2016; Romee et al., 2012), and we confirmed increased expression of IFN-*γ* in these cells by cytokine re-stimulation which is accorded with previous reports (supplementary figure Fig. S2). Compared to freshly isolated, or non-stimulated control NK cells, we found that percentage of CD56^*bright*^ CD62L^+^ cells were higher in CIML NK cells (Fig 1A). Adhesion molecule CD62L expression is expressed on all resting CD56^*bright*^ NK cell, but within CD56^*dim*^ cells, only immature CD57^*-*^ subset expressed it, then decreased during cell maturation (Juelke et al., 2010; Moretta, 2010). In addition to CD62L, CD94/NKG2A^+^ cells, but not NKG2C^+^ cells were also increased (Fig 1B) in CIML NK cells. NKG2A is expressed on the most of resting CD56^*bright*^ NK cell, thus these results show that CIML NK cells are closely related with highly cytokine producing CD56^*bright*^ CD62L^+^ immature NK cells. On the other hand, NKp80 expression level, and terminal maturation marker CD57, and CD16 (Fc*γ*RIIIa) positive cells are decreased in CIML NK cells. There were no significant differences between control and CIML NK cells for pan-KIRs (KIR2Ds and KIR3DS1/L1), NKG2D, and DNAM-1 expression. Intensity of intracellular cytotoxic granules perforin and granzyme B is higher in CIML NK cells. As phenotype of CIML NK cells are similar to CD56^*bright*^ CD16^*-*^ immature NK cells, thus, we hypothesized that CIML NK cells are derived from CD56^*bright*^ immature NK cells.

**Figure 1.**
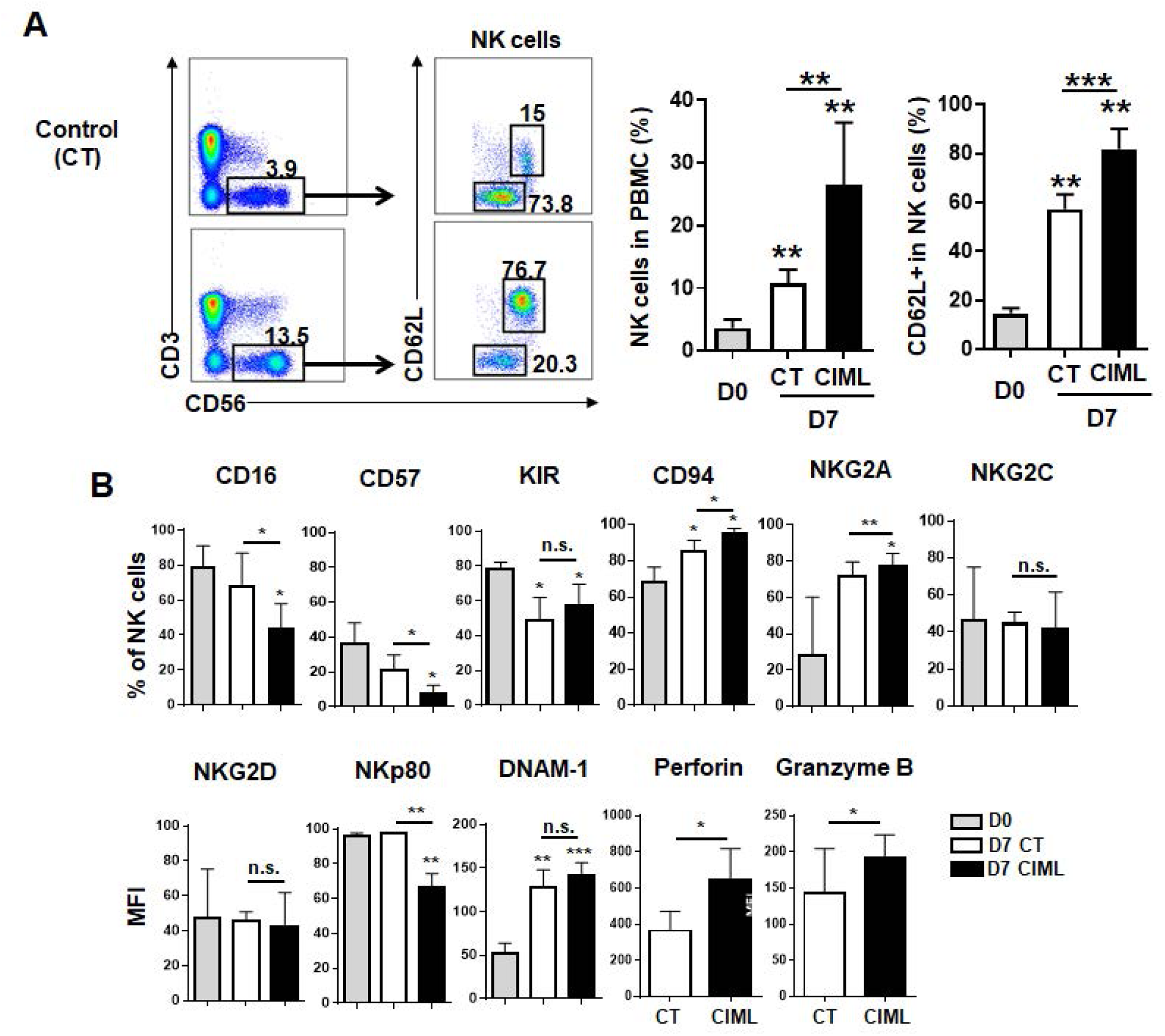
CD56brightCD62L+ cell subsets were enriched in human cytokine induced memory-like NK cells. Phenotype of resting (D0), control and CIML NK cells (D7) were analyzed by flow cytometry. (A) NK cells were gated in CD3- CD56+ cells (left dot plots) then CD56 and CD62L expression on NK cells were showed in right dot plots. FACS plots from one donor are representatively shown from five donors for NK cells (bar graphs). (B) The bar graphs demonstrate that the different expression of cell surface marker between control CIML NK cells (n=4 except for n=3 for DNAM-1, n=5 for granzyme B, and perforin). Statistical significance of all graphs determined with paired T-test. *P<0.05, **P<0.01, and ***P<0.001.

### CD62L+ cytokine-induced memory-like NK cells are highly proliferated CD56bright CD16- CD62L+ immature cells

Within resting NK cells, CD62L^+^ CD57^*-*^ cells have higher proliferative potential compared to CD62L^*-*^ CD57^+^ cells (Bjorkstrom et al., 2010; Juelke et al., 2010; Lopez-Verges et al., 2010). Although low dose IL-15 (1 ng/ml) which others and here we used, is not sufficient to induce robust NK cell proliferation, as shown Fig 2A, CD62L^+^ NK cells highly proliferated after overnight stimulation with CIML-making cytokines. In addition, based on CD56 intensity and CD62L expression, only CD56^*bright*^ CD62L^+^ cells proliferated with low dose IL-15 regardless of cytokine stimulation (Fig 2B). Taken together, increased portion of the CD56^*bright*^ CD62L^+^ cell subset in CIML NK cells is the result of cell expansion after cytokine stimulation. CD62L is also expressed on the immature CD56^*dim*^ NK cell subset (Juelke et al., 2010), and is shed after NK cell activation and proliferation (Romee et al., 2013), thus to clarify selective expansion of CD56^*bright*^ CD62L^+^ cells, next we purified resting NK cell subsets into three populations based on expression of CD56 density, and CD16 and CD62L expression; 1) CD56^*bright*^ CD16^*-*^ (all are CD62L^+^) immature cells, 2) CD56^*dim*^ CD16^+^ CD62L^+^ less mature cells, and 3) CD56^*dim*^ CD16^+^CD62L^*-*^ mature cells (supplementary figure FigS3). These three NK cell subsets were stimulated or rested for overnight, and further cultured for 7 days in the same way above. Whereas about 70% of non-stimulated CD56^*bright*^ CD16^*-*^ CD62L^+^ cells lost CD62L expression, the CIML NK cells maintain CD62L^+^ phenotype (Fig 2C and D). In contrast, regardless of cytokine stimulation, about 30% of CD56^*dim*^ CD62L^+^ cells lost CD62L expression. CD62L was not acquired on the CD56^*dim*^ CD16^+^ CD62L^*-*^ cells. There were no significant changes of CD16, NKG2A, and NKG2C expression by cytokine stimulation within all three subsets (supplementary figure FigS4). CIML NK cells from the CD56^*bright*^ CD16^*-*^ CD62L^+^ subset did not acquire CD16 and NKG2C, but KIR expression was induced compared to control cells (see below). Thus, dominant population with CIML NK cells is derived from CD56^*bright*^ CD16^*-*^ CD62L^+^ NK cells, but not other cell subsets, due to the higher proliferation capacity in response to cytokines. Actually CD56^*bright*^ cells have higher cytokine responsibility (Juelke et al., 2010) and express other cytokine receptors such as IL-7R*α* and SCFR, which are not expressed on CD56^*dim*^ cells (data not shown). In fact IL-18R*α* expression is much higher on CD56^*bright*^ CD16^*-*^ cells than other NK cell subsets, however, a IL-12 receptor IL-12R*β* 1 level was lower (supplementary figure FigS5). As IL-12, rather than IL-18, is a key cytokine for generating CIML (see below), this is not because of cytokine receptor expression levels.

**Figure 2.**
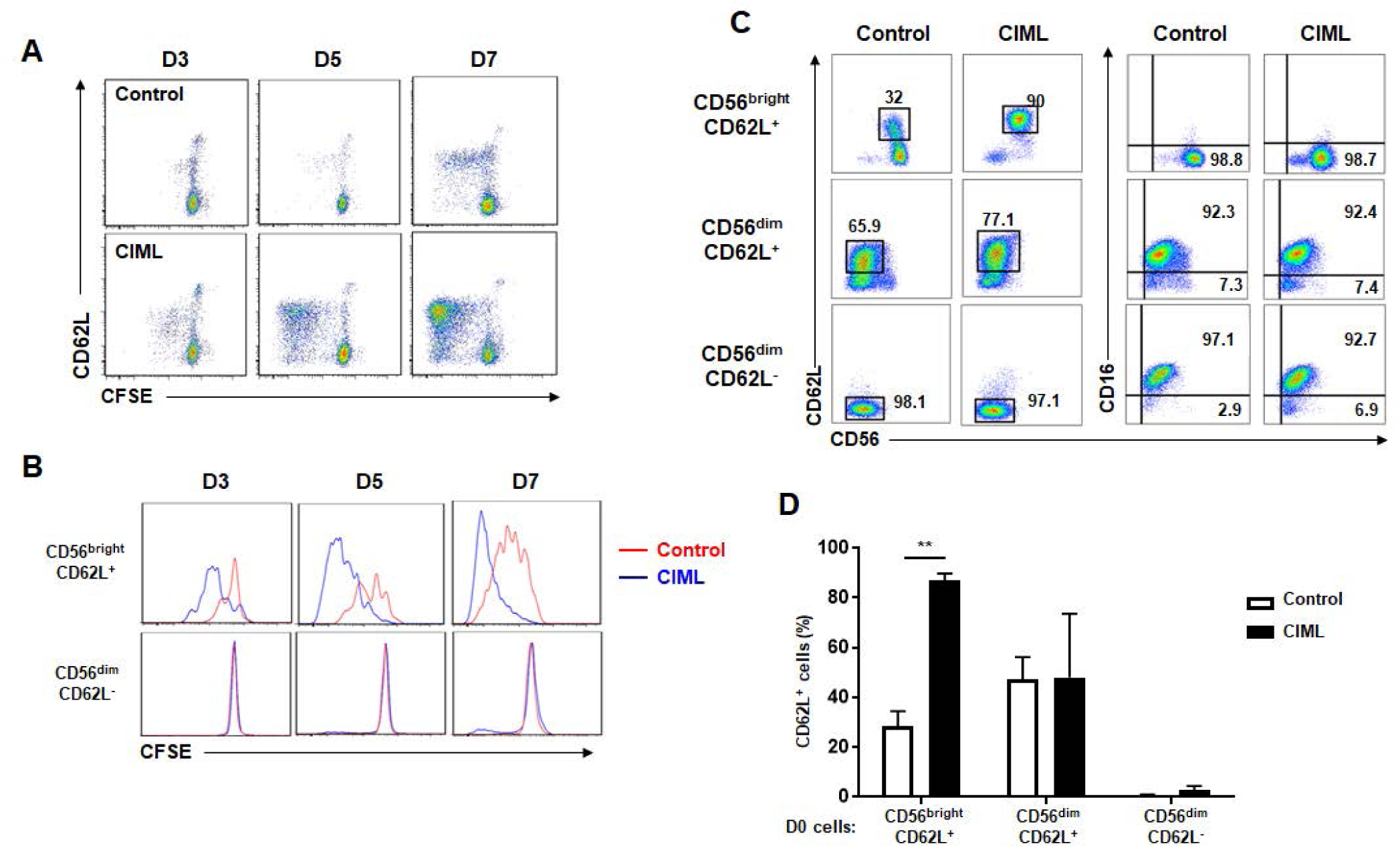
CD56bright CD62L+ NK cell subset was selectively proliferated after cytokine pre-activation. Control and overnight cytokine-activated NK cells were harvest, washed with fresh media, and labeled with 5 M CFSE then further cultured for 7 days in the presence of IL-15 (1ng/ml). On 3, 5, and 7 days after stimulation, cells were stained with CD62L, CD3, and CD56. (A CD62L expression and CFSE were showed gated on NK cells (one representative data shown from more than five independent experiments). (B) Histograms present comparison of cell proliferation in CD56bright CD62L+ and CD56dim CD62L-subsets of control (red) and CIML (blue) NK cells. (C and D) Fresh resting NK cells were FACS-sorted into three populations based on their expression of CD56(bright/dim), CD16, and CD62L. Sorted NK subsets were not stimulated (control) or stimulated (CIML) and further cultured for 7 days. (C) FACS plots are representative data from one donor. (D) The bar graph presents CD62L expression on each sorted group (n=3). Statistical significance analyzed by paired T-test (**P<0.01).

### Eomesodermin is highly expressed on CD62+ CIML NK cells

We confirmed enhanced IFN-*γ* production on CIML NK cells by cytokine re-stimulation. However, there were no difference of degranulation between control and CIML cells (supplementary figure FigS2) in spite of higher cytotoxic granule expression. So we measured and compared the expression of transcription factors which regulate NK cell differentiation, cytokine production, and cytotoxicity. Eomes (eomesodermin) expressing cells were increased on CIML NK cells compared to fresh isolated or control NK cells (Fig 3A-C). Within the both CD62L^+*/-*^ subsets, Eomes^*-*^ cells were disappeared after cytokine stimulation, but Eomes expression is much higher on CD62L^+^ cells. In contrast, T-bet expression is down-regulated, compared to fresh NK cells, but no differences of % positive or intensity were seen on CD62L^+*/-*^ or cytokine treatment^+*/-*^ populations (Fig 3D-F). We also saw similar results from PLZF expression, whereas Helios expression was extremely heterogeneous, and no relevance to CD62L expression or cytokine stimulation (both data not shown).

**Figure 3.**
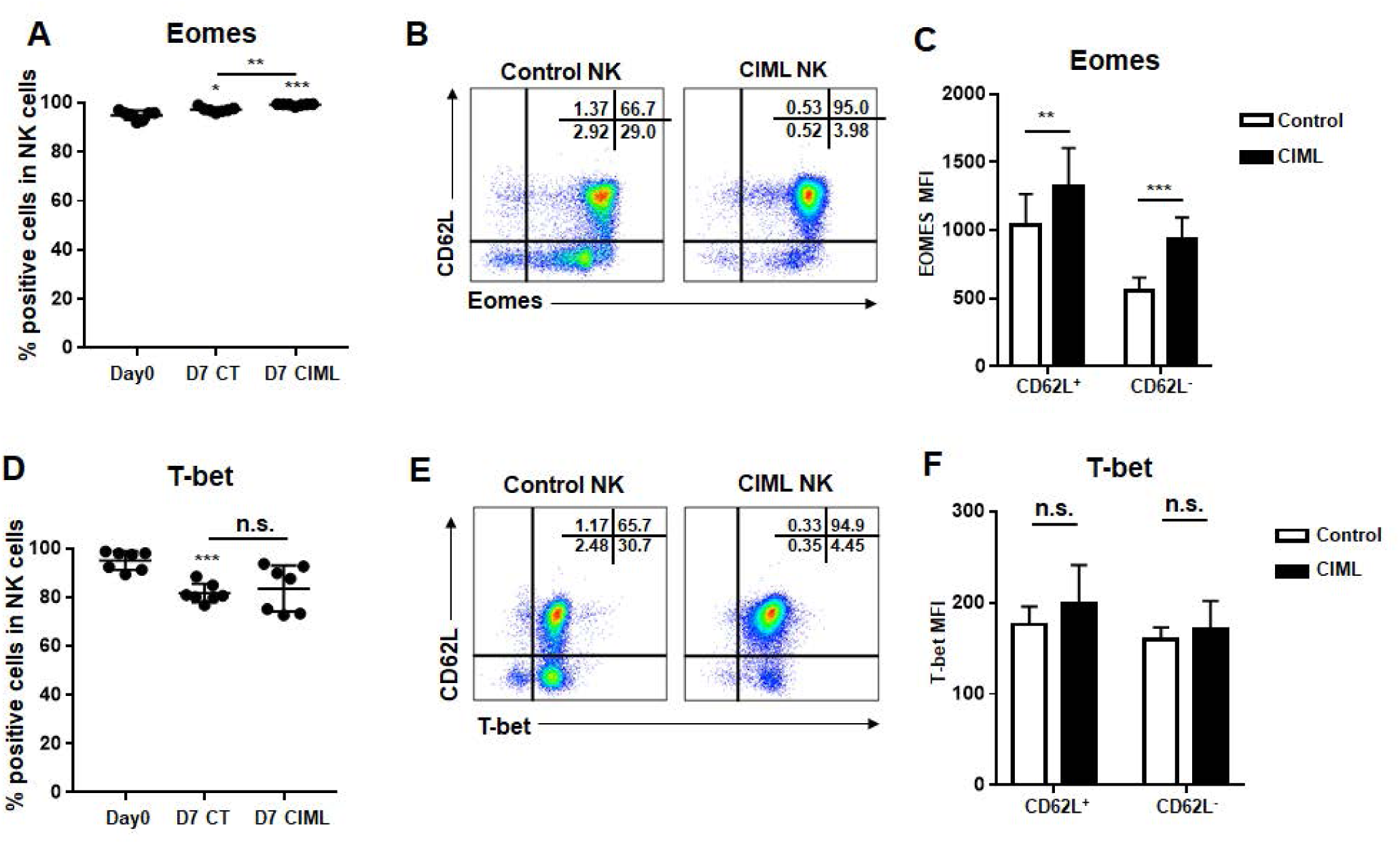
Eomes expression is increased on CIML CD56bright CD62L+ NK cells. Eomes and T-bet expression level were intracellularly measured on resting (D0), control, and CIML NK cells (D7). The graph shows expression level of Eomes (A and B), T-bet (D and E) positive cells in NK cells (n=7). Transcription factor expression is also shown on CD62L+ or CD62L-, control or CIML NK cell populations (n=7). (C and F). FACS plots demonstrate data from representative one donor. Statistical significance analyzed by paired T-test. *P<0.05, **P<0.01, ***P<0.001, and n.s. (not significant) P>0.05

### Cytotoxic CIML NK cells acquired KIR expression, but a negative checkpoint protein NKG2A was also induced and inhibits CIML NK cell functions

CD94 protein can make a heterodimer complex with either inhibitory receptor NKG2A or activating receptor NKG2C, but not both. Without CD94 expression, NKG2A neither NKG2C was not expressed. Therefore, NK cells can be classified into three subsets: CD94/NKG2A^+^ NKG2C^*-*^, CD94/NKG2C^+^ NKG2A^*-*^, and triple negative cells. Whereas terminally maturated CD94/NKG2C^+^ NKG2A^*-*^ cells express KIR, CD56^*bright*^ NK cells (which give rise to most of CIML cells) are CD94/NKG2A^+^ NKG2C^*-*^ and mostly KIR^*-*^ (Bjorkstrom et al., 2010). In addition to CD62L, as shown in Fig 4A and B, we found that NKG2A-expressing NK cells were increased within CIML NK cells (74.8 ± 17.5% compared to 44.2 ± 16.1% of control NK cells, n=5). More CIML NKG2A^+^ cells co-expressed CD62L (84.7 ± 8.8% compared to 49.4 ± 19.0% of control NK cells, n=5). NKG2A^+^ CD62L^+^ CIML NK cells are mainly derived from CD56^*bright*^ CD16^*-*^ cells, and sustain an immature phenotype, but we found that they acquired KIR expression (Fig 4B), which is associated with enhanced cytotoxicity. Using CD107a degranulation assay against MHC I^*-*^- 721.221 target cells, we found that KIR^+^ NKG2A^+^ CIML NK cells, but not control NK cells, showed increased degranulation (Fig 4C). Although more NKG2A^+^ cells were found within CD107^+^ CIML NK cells against K562 and 721.221 target cells, degranulation of NKG2A^+^ CIML NK cells was decreased against HLA-E-transduced 721.221 cells (721.221-AEH) (Fig 4D). Most of NKG2A^+^ CIML NK cells do not co-express activating receptor NKG2C which showed enhanced degranulation to 721.221-AEH cells (Fig 4E). NKG2A blockade restored decreased degranulation of CIML NK cells against 721.221-AEH cells (Fig 4F). Therefore, NKG2A is a major checkpoint molecule in CIML NK cells especially for HLA-E^+^ tumor.

**Figure 4.**
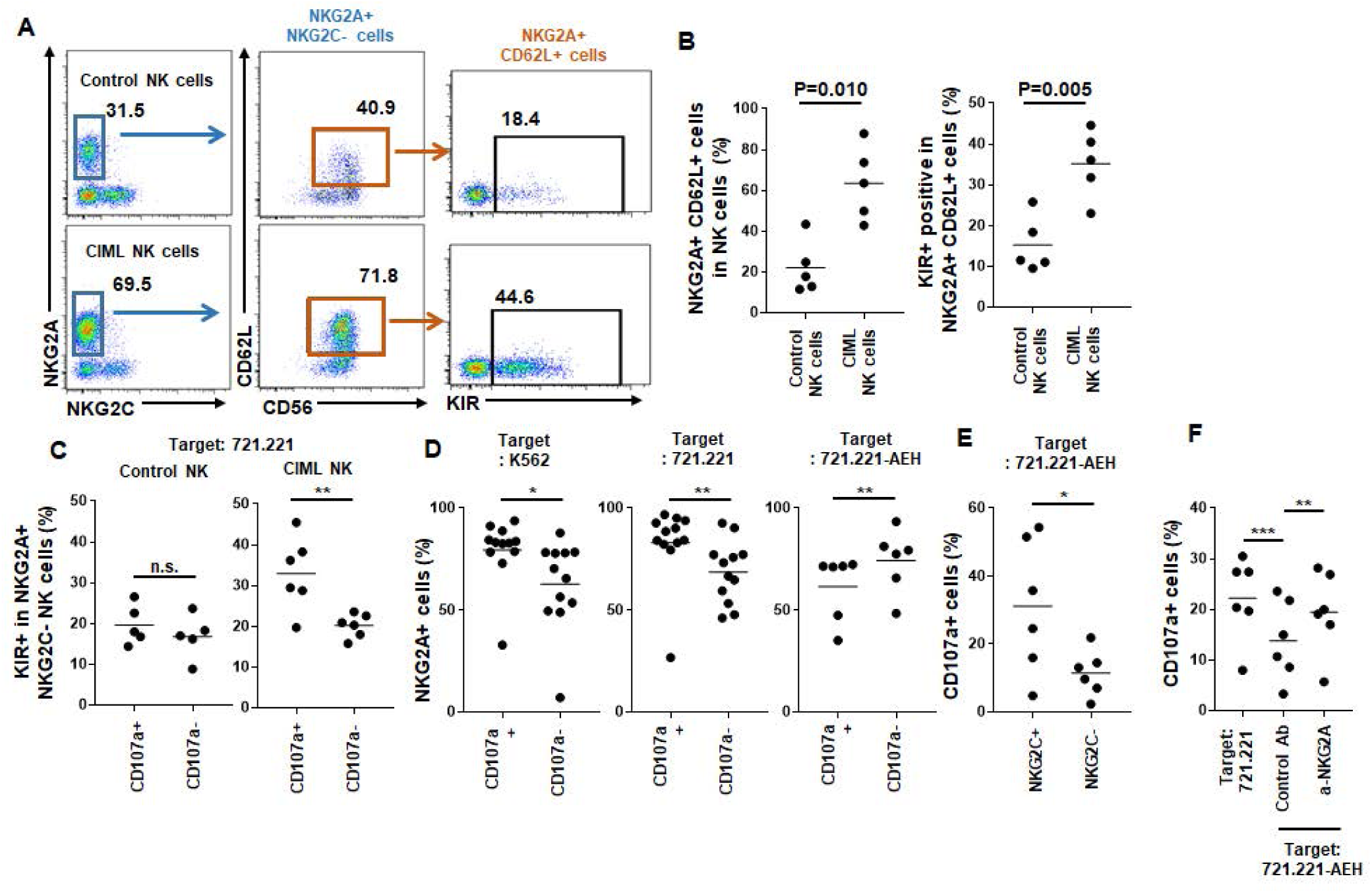
CD62L+ CIML NK cells express NKG2A and KIR (A and B) On day 7, CD62L and KIR expression were analyzed in NKG2A+ control and CIML NK cells (n=5). (C-F) Cytotoxicity of NK cell subsets were analyzed using CD107a degranulation assay co-culture with MHC-I- HLA-E- K562 and 721.221 cells or HLA-E-transduced 721.221-AEH cells. (C) KIR-expressing cells were measured in CD107a+ or CD107a- populations on NKG2A+ NKG2C- control (left, n=5) or CIML NK cells (right, n=6). Percentages of NKG2A+ cells were also shown measured and compared in CD107a+ or CD107a-populations after co-culturing with K562 (D), 721.221 (E), and 721.221-AEH (F) target cells. (E) Degranulation against 721.221-AEH target cells was also measured in NKG2C+ or NKG2C-cells in CIML NK cells. (F) Total CD107a+ CIML NK cells were measured after co-cultured with 721.221 (left), or 721.221-AEH cells in the presence of NKG2A blocking antibody (right) or control mouse IgG (center). Statistical significance of all graphs determined with paired T-test. *P<0.05, **P<0.01.

### LAG-3 is highly expressed in NKG2A- NKG2C+ CIML NK cells

The checkpoint inhibitor PD-1 is also expressed in NK cells in cancer patients, but at lower level compared to other immune cells (Beldi-Ferchiou et al., 2016; MacFarlane et al., 2014). Monalizumab, a blocking antibody CD94/NKG2A-mediated inhibitory signaling is promising target for NK cells especially for CIML NK cells as shown above. Next we investigated other targetable inhibitory receptors on NKG2A^*-*^ CIML cells as terminally maturated cytotoxic NK cells including NKG2C^+^ adaptive NK cells do not express NKG2A. We found that LAG-3, which is expressed on tumor-infiltrating T cells, is expressed on CIML cells from some donors, but not resting or control NK cells. Interestingly, LAG-3 is mainly expressed on the NKG2C^+^ populations only within CIML cells (Fig 5A). LAG-3 expression was detected on day 3 after cytokine stimulation, picked on day 7, then decreased on day 10 (Fig 5B). As LAG-3 is expressed on exhausted T cells, we speculated that mature, exhausted NK cells may express it. LAG-3 expression was further measured within CIML NK cell subsets (when LAG-3 is expressed more than 5% of NK cells) based on the NKG2A, NKG2C, and KIR expression. As shown in Figure 5C and D, LAG-3 expression is increased through their maturation status; NKG2A^+^ KIR^*-*^ (immature: 6.4 ± 3.6%), NKG2A^+^ KIR^+^ (mature: 17.6 ± 12.4%), and NKG2C^+^ KIR^+^ (terminally matured: 54.7 ± 25.4%). Whereas exogenous IL-18 alone, type I interferon, or high-dose IL-2 or IL-15 did not induce LAG-3 expression on NK cells (data not shown), IL-12 was a main inducer of LAG-3, and IL-18 further enhanced it (Fig 5E). In summary, CIML-inducing short term *in vitro* cytokine activation, by IL-12 and IL-18, induces selective expansion of CD56^*bright*^ NKG2A^+^ CD62L^+^ NK cell with acquisition of KIR expression. Also it induces LAG-3 expression in NKG2C^+^ KIR^+^ and NKG2A^+^ KIR^+^ mature, minor NK cell populations. Therefore, dual-blocking NKG2A and LAG-3 can be considered for two CIML NK cell populations.

**Figure 5.**
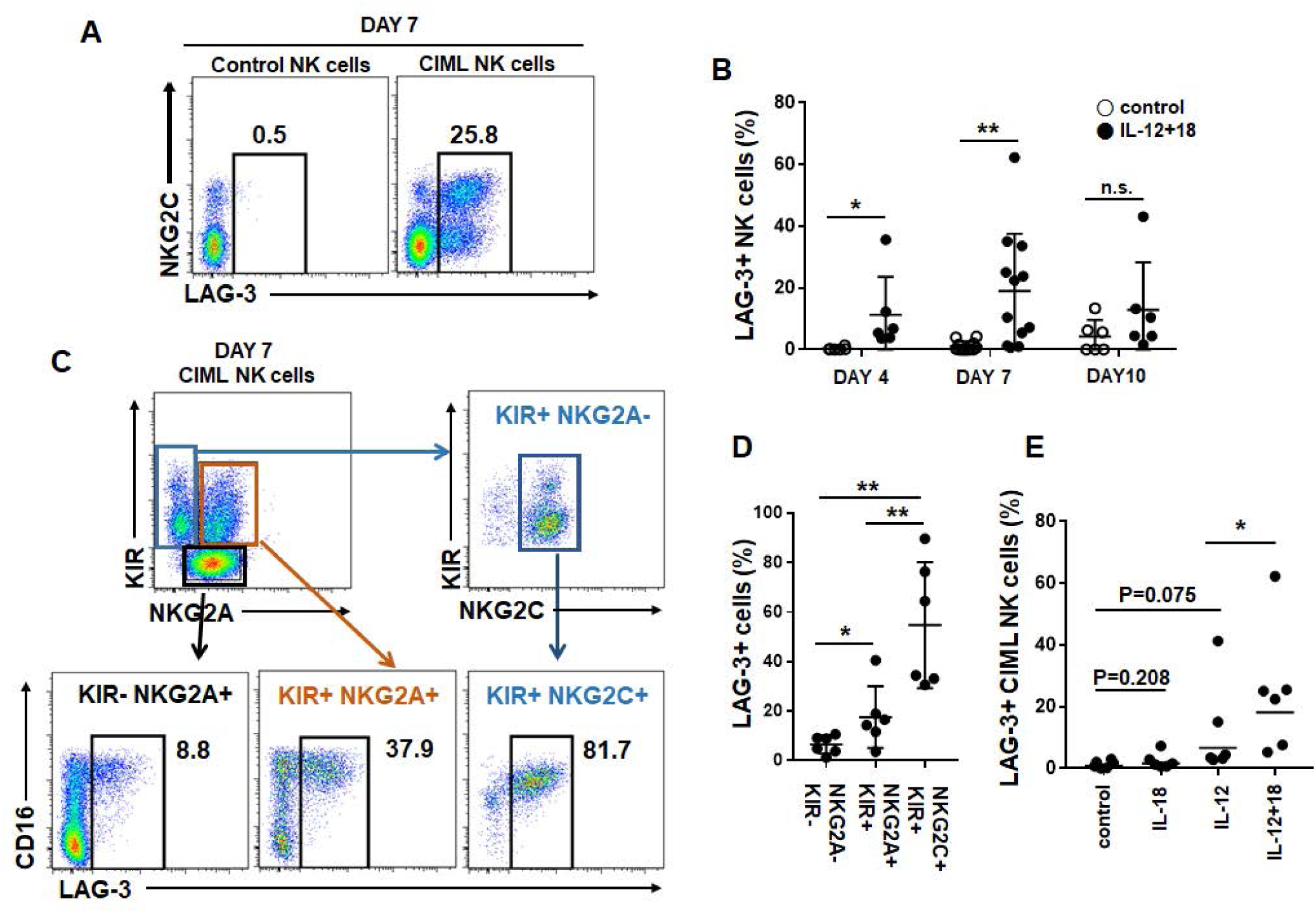
LAG-3 expression is induced on NKG2C+ CIML NK cells LAG-3 expression was analyzed on control (left) and CIML NK cells (right) on day 7 (A), and also on day 3 and 10 (B). (C and D) As LAG-3 is mainly expressed on NKG2C+ CIML NK cells, LAG-3 is further analyzed on CIML NK cell subsets based on NKG2A, NKG2C, and KIR expression. (E) In some experiments, NK cells were stimulated with IL-12 alone, IL-18 alone, or IL-12 and IL-18 (CIML), then LAG-3 expression is analyzed on day 7 (n=6). Control NK cells were not stimulated with IL-12 or IL-18. Statistical significance of all graphs determined with paired T-test. *P<0.05, **P<0.01.

## Discussion

In 2009, Cooper et al., reported the presence of immunological memory in murine NK cells for the first time. 1-3 weeks after short-term *in vitro* cytokine-stimulation, NK cells do not express IFN-*γ* constitutively, but in response to subsequent simulation, they express significant level of IFN-*γ* (Cooper et al., 2009). This finding was also confirmed in human NK cells (Leong et al., 2014; Romee et al., 2012), and CIML NK cell therapy clinical trials are ongoing (Romee et al., 2016). In other hand, unique human NK cell population lacking an adaptor protein Fc*ε*RI*γ* was reported (Zhang, Scott, Hwang, & Kim, 2013). Self MHC I-recognizing KIR, NKG2C, and maturation marker CD57 are expressed on the cells, and their DNA methylation profile is similar to effector CD8+ T cells (Lee et al., 2015; Schlums et al., 2015). Without several signal transduction protein expressions, adaptive NK cells have superior cytotoxicity. CMV infection is associated with i*n vivo* generation of adaptive NK cells (Beziat et al., 2013), and leukemia patients who receive hematopoietic stem cell transplantation with CMV-positive donors have lower relapse rate with increased NKG2C^+^ CD57^+^ cell numbers in the blood (Foley, Cooley, Verneris, Curtsinger, et al., 2012; Foley, Cooley, Verneris, Pitt, et al., 2012). Although induction of adaptive NK cells by MCMV and HCMV is not antigen-specific manner, murine m155 and human HLA-E is critical for clonal expansion of NKG2C^+^ cells (Rolle et al., 2014; Sun, Beilke, & Lanier, 2009). Although IL-12 and IL-18 also have important roles in CMV infection and NK cell activation (Madera & Sun, 2015; Rolle et al., 2014), cytokine-induced memory-like NK cells differ from CMV-driven adaptive NK cells. Here we show that CIML NK cells are mainly cytokine-driven CD56^*bright*^ immature NK cells and have similar features with immature NK cells. CD56^*bright*^ CD16^*-*^ NK cells have higher cytokine responsibility, express abundant IFN-*γ*, have a long lifespan (Romagnani et al., 2007), so they can highly proliferate in response to IL-12 and IL-18 without high dose of IL-15.

Based on discovery of NK cell memory, NK cell immunotherapy strategies have been changed. Adaptive transfer of CIML NK cells have several advantages which are long lifespan and higher cytokine responsibility as *in vivo* persistency of transferred NK cells is short, and administration of cytokines such as IL-2 or IL-15 have cytotoxicity. Importantly, CIML NK cell sustain CD62L expression which is usually down-regulated during *ex vivo* NK cell activation (Romee et al., 2013). As CD62L is an important homing receptor of NK cells to migrate into BM and secondary lymphoid tissues (Olson, Zeiser, Beilhack, Goldman, & Negrin, 2009; Persson & Chambers, 2010), sustain CD62L expression on once activated NK cells may improve *in vivo* persistency after adaptive transfer. In other hand, adaptive NK cell-based immunotherapies have also been tried. These KIR^+^ NKG2C^+^ CD57^+^ highly cytotoxic and mature NK cells are selectively expanded from CMV+ donor NK cell. Mature, adaptive NK cells do not proliferate well, instead immature NK cells were maturated using a pharmacological inhibitor of GSK3*β* kinase (Cichocki et al., 2017). Otherwise, NKG2C^+^ NK cells are selectively activated and expanded using human HLA-E expressing feeder cells (Liu et al., 2017). In contrast to adaptive NK cells, CIML NK cells dominantly express NKG2A which is an inhibitory receptor for HLA-E. Although enhanced cytotoxicity of CIML NK cells have shown, however, widely used K562 and 721.221 target cells used for measuring NK cell cytotoxicity do not express either MHC I and HLA-E. Thus for the patients who have higher HLA-E expression level in the tumor tissues, use of NKG2A blocking antibody should be considered for CIML NK cell therapy. LAG-3 is also induced on CIML NK cells especially on the further maturated cells (NKG2C^+^ KIR^+^ and NKG2A^+^ KIR^+^ cells), thus blocking LAG-3 also should be considered for the tumors expressing MHC II, the inhibitory ligand of LAG-3. MHC II is constitutively expressed normal and malignant immune cells, and also can be induced in solid tumor cells by tumor-infiltrating activated immune cell-derived IFN-*γ* in tumor microenvironment.

In summary, here we showed that short term *in vitro* cytokine stimulation actually changes dominance of NK cell subsets rather than generating memory-like functions. Also, we showed that dominantly expanded NK cells express NKG2A, whereas minor NKG2C^+^ cells acquired LAG-3 expression. Therefore, blockade of these molecules should be considered for use of CIML NK cells.

## Methods and Materials

### Cell isolation and culture

Peripheral blood mononuclear cells (PBMCs) from healthy donors were separated using Ficoll-Hypaque density gradient from de-identified leukoreduction system chambers (LRS) chambers after routine healthy donor plateletpheresis procedures at Seoul National University Hospital. The institutional review board of SNUH reviewed and exempted this study. Freshly isolated or thawed frozen PBMCs were seeded at 3-4E+06/ml in the media supplemented with row dose rhIL-15 (1 ng/ml, prospec) as a survival factor. To generate CIML NK cells, cells were stimulated with rhIL-12 (10 ng/ml, peprotech) and rhIL-18 (50 ng/ml, R&D systems) for overnight. Control cells were cultured with row dose IL-15 only. Cells were harvested, washed cultured in fresh media with IL-15 (1 ng/ml) for 7-14 days. During the culture, half of the media was replaced with fresh media containing IL-15 twice a week. Cells were cultured in DMEM: HAM’s F12 (at 2:1 ratio) media supplemented with 10% heat inactivated human AB serum (Gemini Bio-Products), 10 *µ*g/ml gentamicin, 25 *µ*M*β*-mercaptoethanol (Gibco), 5 ng/ml sodium selenite, 50 *µ*M ethanolamine, 20 *µ*g/ml ascorbic acid, and 1 mM nicotinamide (Sigma).

### Antibodies and flow cytometry

Phenotype of NK cells were analyzed by flow cytometry. Florescence-conjugated antibodies against CD3 (clone BW264/56), KIR2D (NKVFS1), KIR3DL1/S1 (REA168), NKG2C (REA205), NKG2A (REA110), CD57 (REA769) and NKp80 (REA845) were from Miltenyi Biotec. Antibodies targeting CD16 (CB16), Helios (22F6), T-bet (4B10), Eomes (WD1928), perforin (dG9), LAG-3 (C9B7W), and IFN-*γ* (4S.B3) were obtained from eBioscience. CD56 (B159), CD62L (DREG-56), CD94 (HP-3D9), NKG2D (1D11), PLZF (R17-809), granzyme B (GB11), and CD107a (H4A3) antibodies were purchased from BD Biosciences. NK cells in PBMCs were gated in CD16^+^ and/or CD56^+^ from CD3^*-*^ population in PBMCs, and dead cells were excluded using viability dye (eBioscience). To detect intracellular proteins, BD Cytofix/Cytoperm Fixation/Permeabilization Solution was used according to manufacturers’ instructions. For transcription factors, eBioscience Foxp3 / Transcription Factor Staining Buffer Set was used. All data were acquired with BD FACSCanto II and analyzed using FlowJo (version 7.6.5., Flowjo).

### NK cell functional assay

To detect cytotoxic NK cell subsets, CD107a degranulation assay was used. K562 cells were obtained from ATCC. 721.221 and 721.221-AEH cells were kindly provided from Dr. Eric Long (National Institutes of Health), and Dr. Dan Geraghty (Fred Hutchinson Cancer Research Center) respectively. Cells were cultured with RPMI1640 supplemented with 10% (for K562) or 15% (for 721.221 and 721.221-AEH) of FBS and gentamycin. NK cells and target cells were co-culture at the ratio of 1:1 for 6 hours in the presence of CD107a antibody and monemsin. (GolgiStop from BD). In some experiments, NK cells were pre-incubated with 10 *µ*g/ml of NKG2A blocking or control antibody (clone Z199 from Beckman Coulter, and mouse IgG from R&D Systems) for 30 min before co-culture. Co-cultured cells were harvested, and stained with CD3, CD16, CD56, NKG2A, NKG2C, and KIR antibodies with dead/live dye. IFN-*γ* expression was measured by flow cytometry (using BD cytofix/cytoperm intracellular staining kit) 6 hours after IL-12 (10 ng/ml) and IL-18 (50 ng/ml) stimulation in the presence of monemsin.

### Cell Proliferation analysis

To track cell proliferation between CD56^*bright*^ CD62L^+^ and CD56^*dim*^ CD62L^*-*^ NK cell populations, overnight-stimulated or control cells were labeled with 5*µ*M CFSE (Thermo Fisher), washed, and cultured. Proliferation was analyzed at day 3, 5, and 7.

### Cell sorting

NK cells were enriched from PBMCs by depleting CD3, CD14, and CD19 positive cells by MACS system (Miltenyi biotec). NK cell subsets were further purified by FACS cell sorting (BD LSR II) based on expression of CD56, CD16, and CD62L in CD3^*-*^ CD56/CD16^+^ NK cells. Purified cells were separated into two groups (CIML and control NK cells), and cultured in the same way above.

### Statistical analysis

Statistical significances were determined by paired or unpaired t-test using GraphPad Prism software version 7.

## Supplementary data: Supplementary figures 1-5

**Figure S1.**
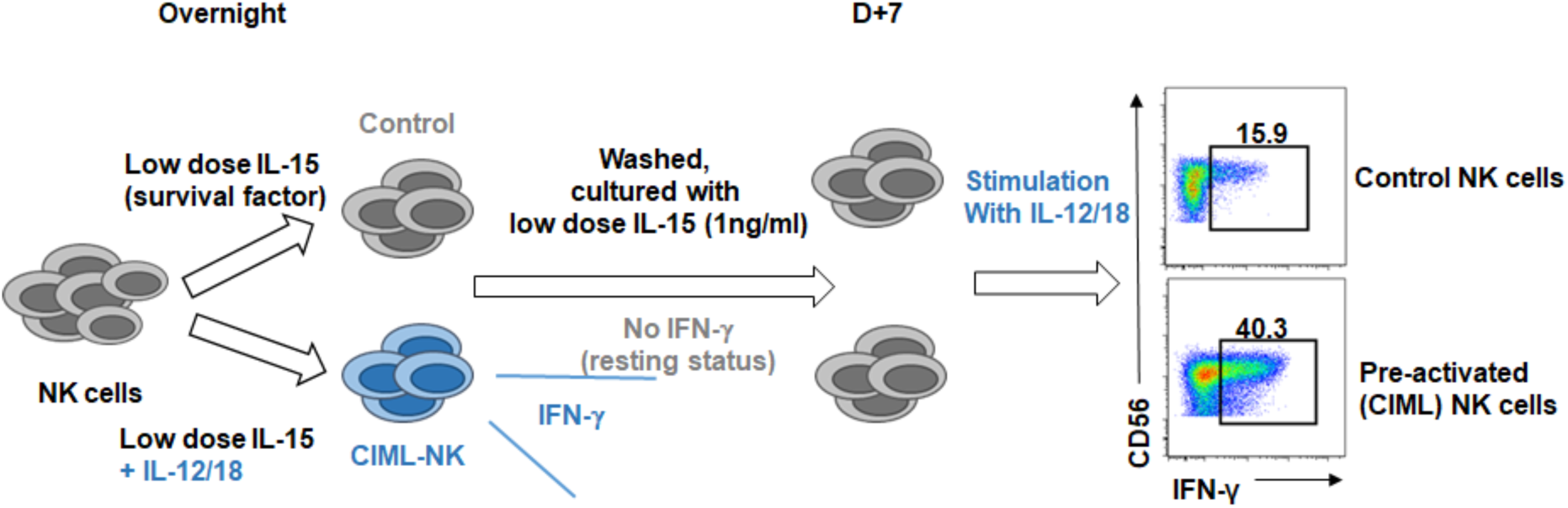
Schema of human cytokine induced memory-like NK cells. Cytokine-induced memory-like NK cells were generated by in vitro stimulated with rhIL-12 (10 ng/ml) and rhIL-18 (50 ng/ml) in the presence of low dose rhIL-15 (1 ng/ml) as a survival factor. Cells were harvested after 16 hours, then washed, and further cultured with fresh media with low dose IL-15.

**Figure S2.**
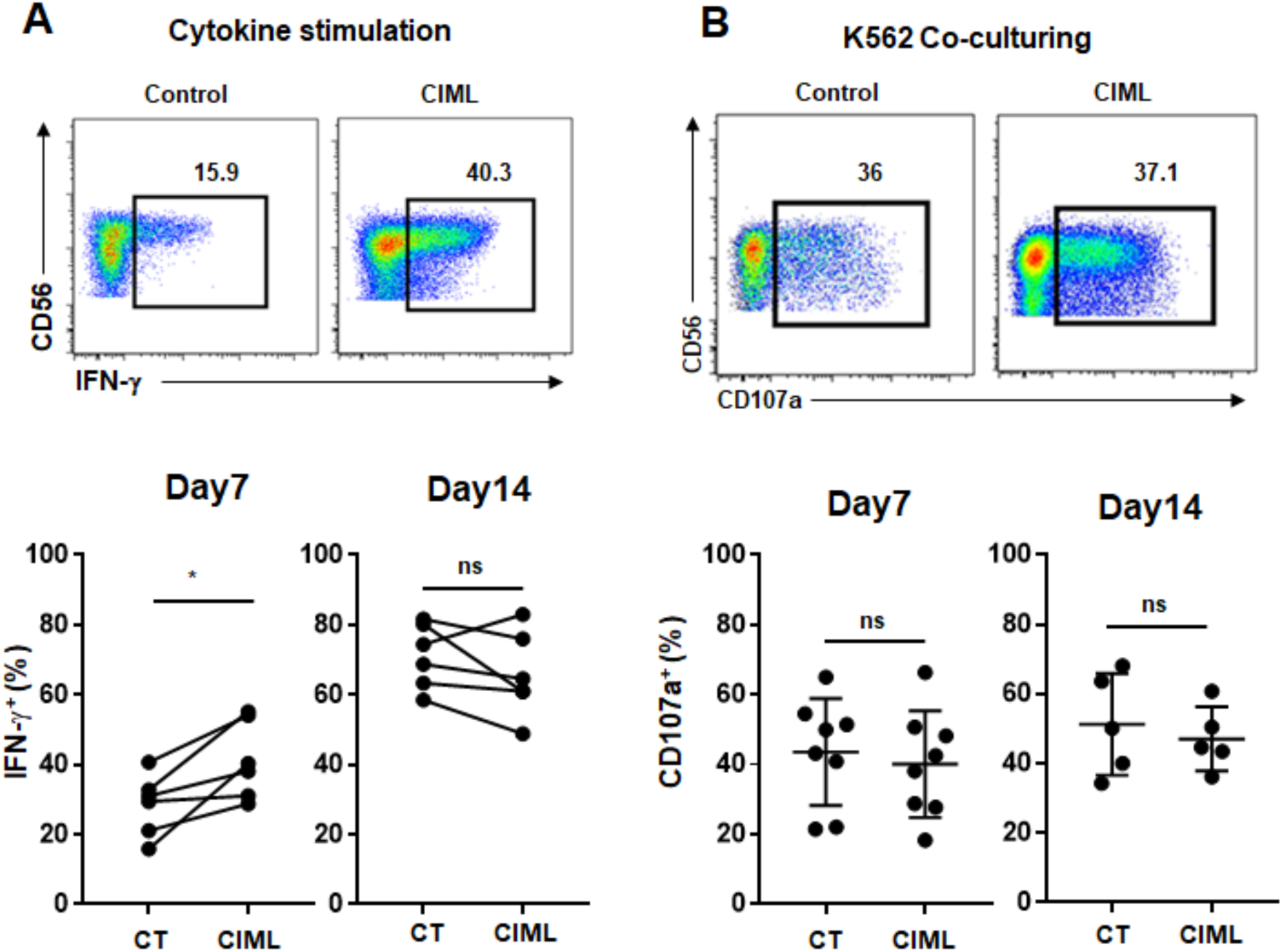
Functional analysis of CIML NK cells. Control or CIML NK cells (on day 7 or 14) were stimulated with IL-12 and IL-18 (A) or K562 co-culturing (B). IFN-γ secretion and CD107a degranulation were measured by flow cytometry.

**Figure S3.**
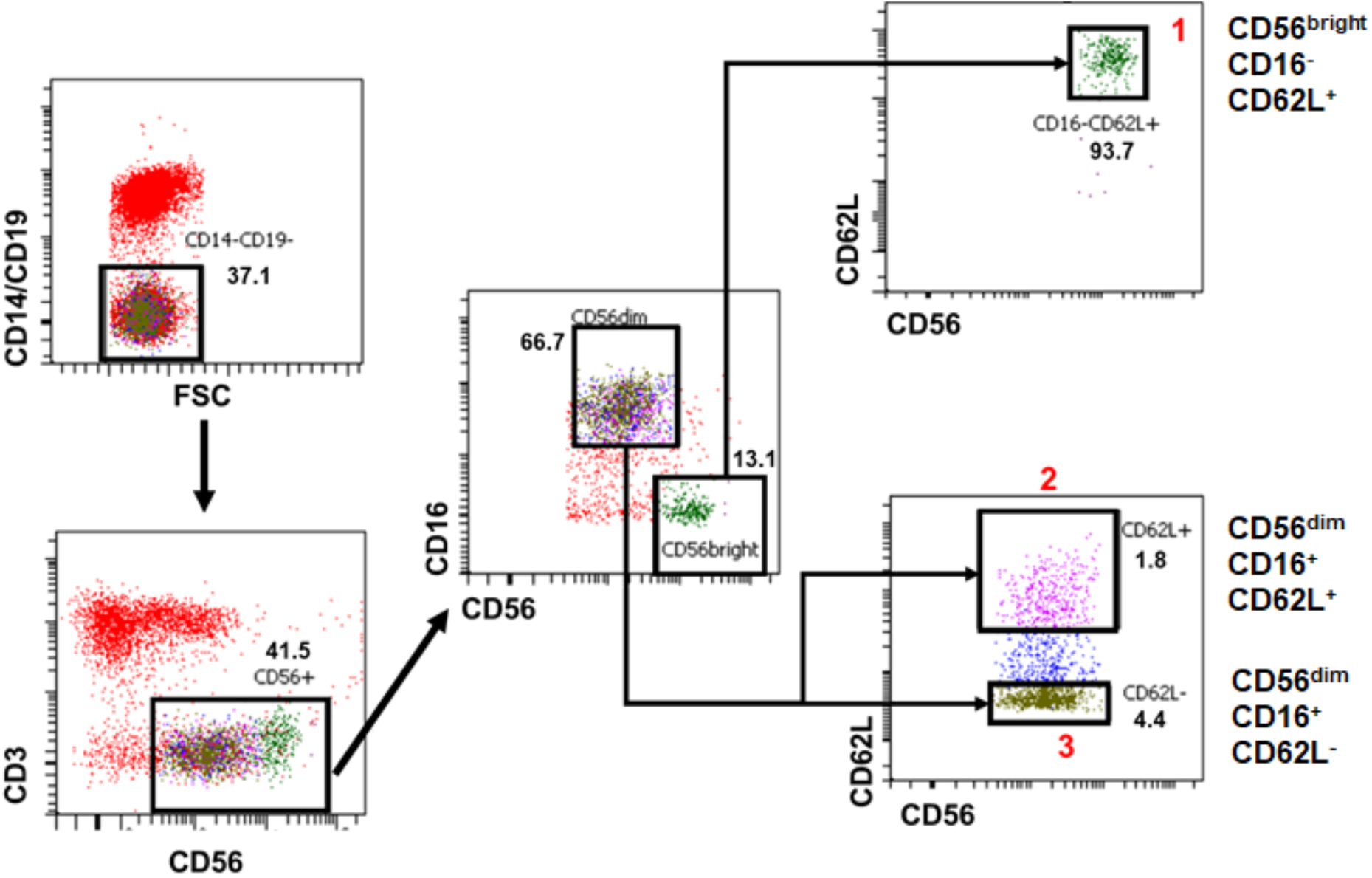
The sorting stage of PBMCs to get three NK cell population. NK cells were enriched from PBMCs and sorted to CD56^bright^CD62L^+^, CD56^dim^CD62L^+^, and CD56^dim^CD62L^−^ NK cell subsets.

**Figure S4.**
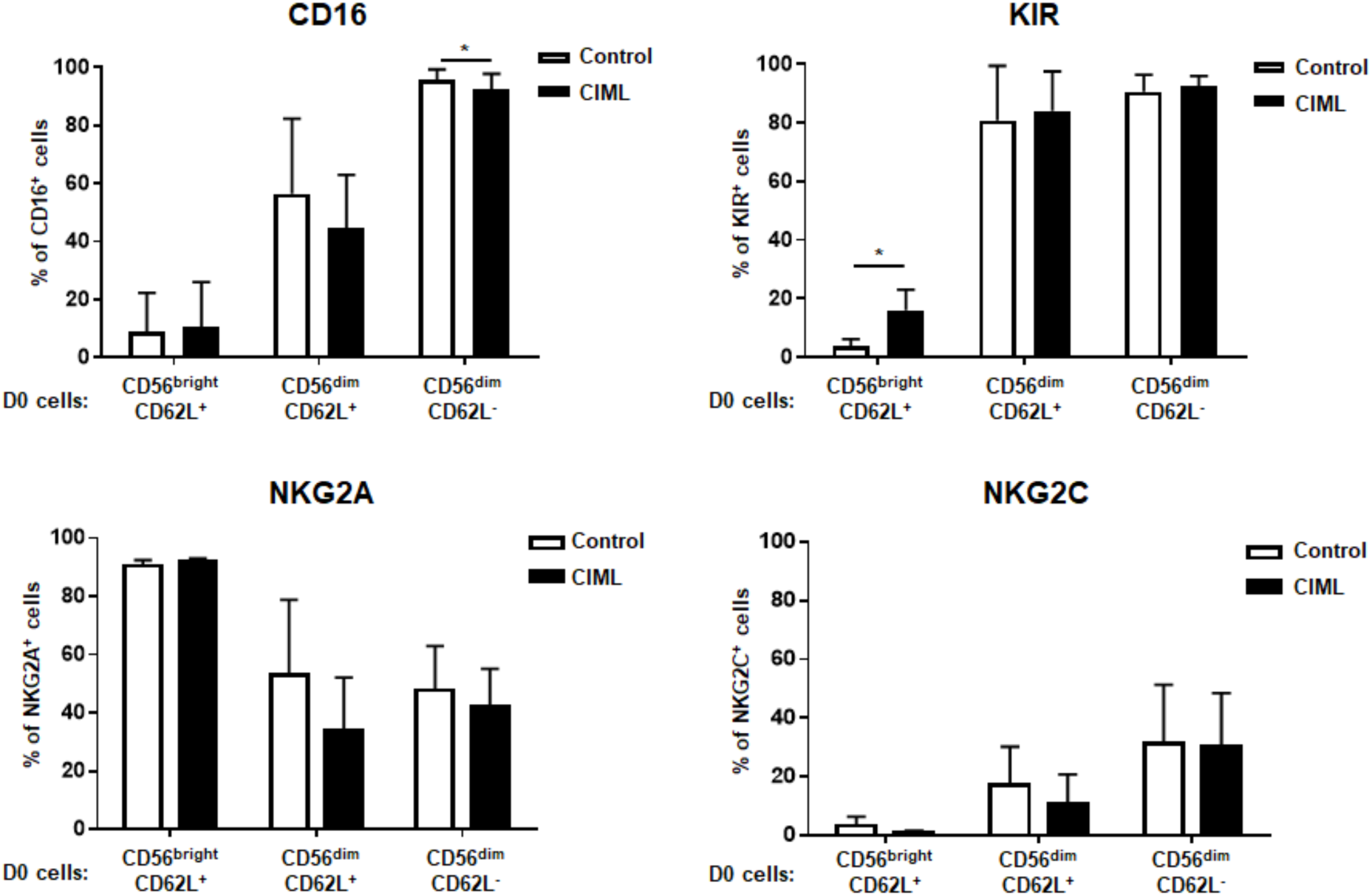
Phenotype of CIML NK cells from three different NK cell populations. FACS-sorted NK cell populations were stimulated with CIML cytokines. 7 days after stimulation, CD16, NKG2A, NKG2C, and KIR expression were analyzed. *P<0.05.

**Figure S5.**
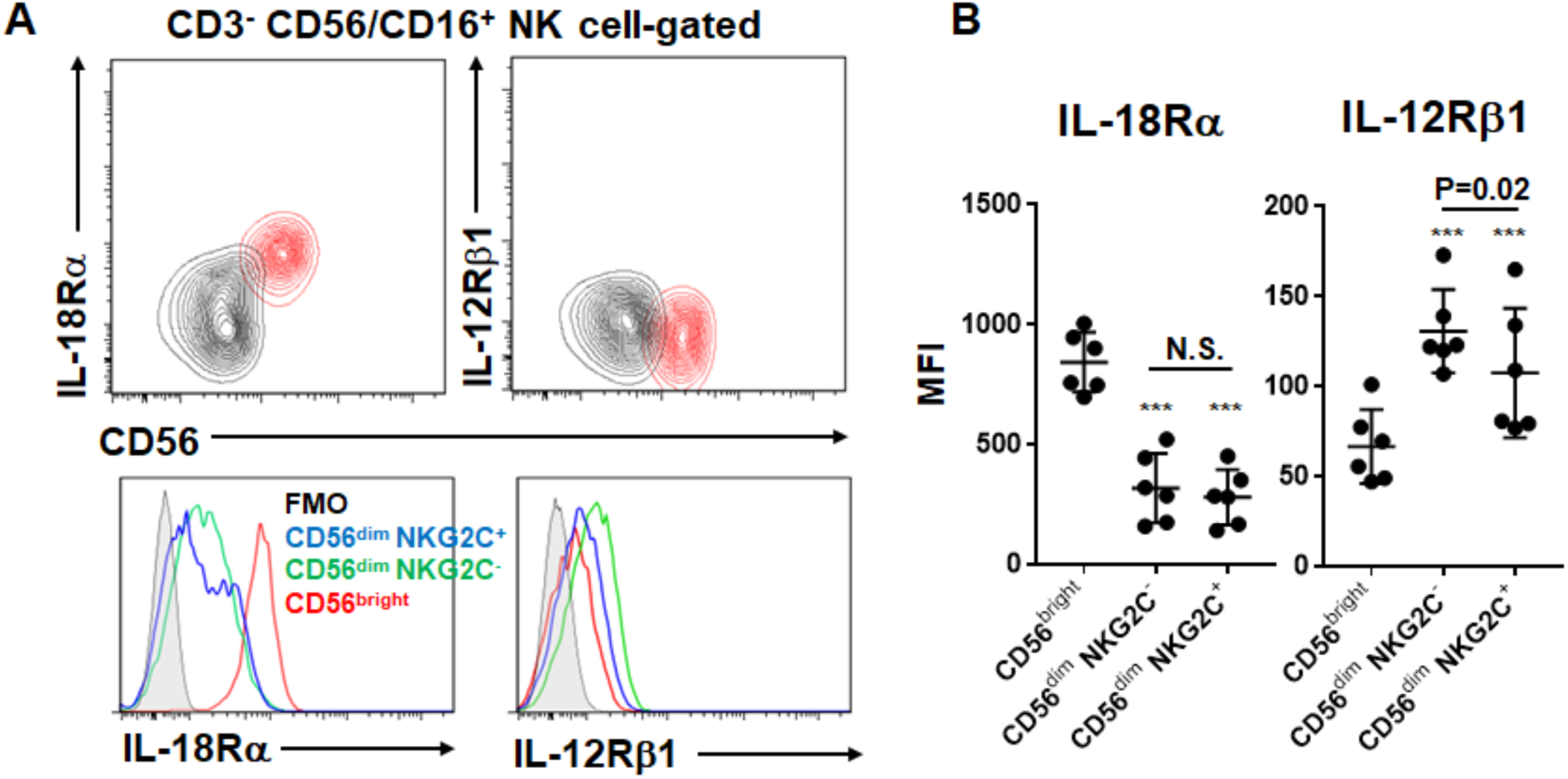
Cytokine receptor expression in NK cell populations. Fresh resting PBMCs were stained with IL-18Rα and IL-12Rβ1 as well as CD3, CD16, CD56, and NKG2C (n=6). Cytokine receptor expression is compared among several NK cell subsets.

